# Changes in subcellular structures and states of Pumilio1 regulate the translation of target *Mad2* and *Cyclin B1* mRNAs

**DOI:** 10.1101/2020.02.25.964668

**Authors:** Natsumi Takei, Yuki Takada, Shohei Kawamura, Atsushi Saitoh, Jenny Bormann, Wai Shan Yuen, John Carroll, Tomoya Kotani

## Abstract

Temporal and spatial control of mRNA translation has emerged as a major mechanism for promoting diverse biological processes. However, the molecular nature of temporal control of translation remains unclear. In oocytes, many mRNAs are deposited as a translationally repressed form and are translated at appropriate timings to promote the progression of meiosis and development. Here, we show that changes in structures and states of the RNA-binding protein Pumilio1 regulate the translation of target mRNAs and progression of oocyte maturation. Pumilio1 was shown to bind to *Mad2* and *Cyclin B1* mRNAs, assemble highly clustered solid-like aggregates, and surround *Mad2* and *Cyclin B1* RNA granules in mouse oocytes. These Pumilio1 aggregates were dissolved by phosphorylation prior to the translational activation of target mRNAs. Stabilization of Pumilio1 aggregates prevented the translational activation of target mRNAs and oocyte maturation. Together, our results provide an aggregation-dissolution model for the temporal and spatial control of translation.

## Introduction

Diverse biological processes including meiosis, embryonic development and neuronal plasticity are promoted by translational activation of dormant mRNAs at appropriate timings and places (Buxbaum et al., 2015; Doyle and Kiebler, 2011; Martin and Ephrussi, 2009; Mendez and Richter, 2001). This temporal control of translation has been most extensively studied in oocyte meiosis. Fully grown vertebrate oocytes are arrested at prophase I of meiosis and accumulate thousands of translationally repressed mRNAs in the cytoplasm (de Moor et al., 2005; Kotani et al., 2017; Masui and Clarke, 1979). In response to specific cues such as hormones, oocytes resume meiosis and are arrested again at metaphase II. This process is termed oocyte maturation and is necessary for oocytes to acquire fertility. For proper progression of oocyte maturation, hundreds of dormant mRNAs are translationally activated in periods specific to distinct mRNAs (Chen et al., 2011). Of these, *Cyclin B1* mRNA, which encodes the regulatory subunit of maturation/M-phase-promoting factor (MPF), is translated in meiosis I, and the newly synthesized Cyclin B1 proteins in this period are prerequisite for the progression of meiosis (Davydenko et al., 2013; Hochegger et al., 2001; Kondo et al., 2001; Kotani and Yamashita, 2002; Ledan et al., 2001; Polanski et al., 1998).

Translational activation of the dormant mRNAs including *Cyclin B1* has been shown to be directed by the cytoplasmic polyadenylation of mRNAs, which is mediated by the cytoplasmic polyadenylation element (CPE) in their 3’ UTR (McGrew et al., 1989; Sheets et al., 1994). The CPE-binding protein (CPEB) functions in both repression and direction of the cytoplasmic polyadenylation (Barkoff et al., 2000; de Moor and Richter, 1999; Gebauer et al., 1994; Tay et al., 2000). Although many dormant mRNAs contain CPEs, they are translated in different periods during oocyte maturation, indicating that there must be additional mechanisms to determine the timings of translational activation of distinct mRNAs. However, the molecular and cellular mechanisms by which translational timings of hundreds of mRNAs are coordinated remain unclear.

Pumilio1 (Pum1) is a sequence-specific RNA-binding protein that belongs to the Pumilio and Fem-3 mRNA-binding factor (PUF) family, which is highly conserved in eukaryotes from yeast to human (Spassov and Jurecic, 2003; Wickens et al., 2002). Pum was identified in *Drosophila* as a protein that is essential for posterior patterning of embryos (Lehmann and Nussleinvolhard, 1987) and it was shown to repress the translation of target mRNAs in a spatially and temporally regulated manner (Asaoka-Taguchi et al., 1999; Murata and Wharton, 1995). In *Xenopus*, zebrafish and mouse oocytes, Pum1 has been shown to bind to *Cyclin B1* mRNA and determine the timing of translational activation of *Cyclin B1* mRNA during oocyte maturation (Kotani et al., 2013; Nakahata et al., 2003; Ota et al., 2011; Pique et al., 2008). Pum1 knockout mice were shown to be viable but defective in spermatogenesis (Chen et al., 2012) and oogenesis (Mak et al., 2016). Pum1-deficient mice also showed neuronal degeneration in the brain through an increase in Ataxin1 protein (Gennarino et al., 2015). Pum1 was shown to target more than one thousand mRNAs in the mouse testis and brain (Chen et al., 2012; Zhang et al., 2017). The amount of proteins synthesized from these Pum1-target mRNAs, but not the amount of mRNAs, was increased in Pum1-deficient mice, indicating that Pum1 represses the translation of target mRNAs (Chen et al., 2012; Zhang et al., 2017). Despite the importance of Pum function in diverse systems, how Pum regulates the translation of target mRNAs remains to be elucidated.

In addition to sequence-specific RNA-binding proteins, we previously demonstrated that formation and disassembly of *Cyclin B1* RNA granules determine the timing of translational activation of mRNA, i.e., granular structures of this mRNA formed in immature, germinal vesicle (GV)-stage oocytes were disassembled at the timing of translational activation of mRNA, and stabilization and dissociation of these granules prevented and accelerated the mRNA translation, respectively (Kotani et al., 2013). Binding of Pum1 was shown to be required for the RNA granule formation, implying that Pum1 regulates the translational timing of target mRNAs through formation and disassembly of granules (Kotani et al., 2013).

P granules are cytoplasmic granules that consist of mRNAs and RNA-binding proteins and have been shown to behave as liquid droplets with a spherical shape in *C. elegance* embryos (Brangwynne et al., 2009). In addition, several RNA-binding proteins that are assembled into stress granules were shown to produce liquid droplets in vitro and in cultured cells (Lin et al., 2015; Molliex et al., 2015). Although phase changes in these liquid droplets into solid-like assemblies have been linked to degenerative diseases (Li et al., 2013; Weber and Brangwynne, 2012), more recent studies have demonstrated the assembly of solid-like substructures within stress granules (Jain et al., 2016; Shiina, 2019; Wheeler et al., 2016), suggesting physiological roles of the solid-like assemblies. However, biological function of phase changes of protein assemblies from liquid to solid states and vice versa remains to be explored.

In this study, we identified *Mad2* mRNA as one of the Pum1-target mRNAs in mouse oocytes and found that *Mad2* and *Cyclin B1* mRNAs were distributed as distinct granules in the cytoplasm. Interestingly, Pum1 was assembled into solid-like aggregates exhibiting highly clustered structures, and these aggregates surrounded *Mad2* and *Cyclin B1* RNA granules. The Pum1 aggregates dissolved in an early period after resumption of meiosis by phosphorylation, resulting in a liquid-like state and translational activation of *Mad2* and *Cyclin B1* mRNAs. These results provide an aggregation-dissolution model, accompanied by phase changes of RNA-binding proteins, for temporal and spatial control of mRNA translation. The results also showed the physiological importance of phase changes of proteins in RNA regulation.

## Results

### Expression of Mad2 is translationally regulated during mouse oocyte maturation

Mad2 has been shown to function as a component of spindle assembly checkpoint proteins to accurately segregate chromosomes in meiosis I of mouse oocytes (Homer et al., 2005). However, how Mad2 is accumulated in oocytes remains unknown. To clarify the mechanism of Mad2 accumulation in meiosis I, we first analyzed the expression of *Mad2* mRNA in mouse oocytes. Although two splicing variants of *Mad*2 mRNA were isolated using purified RNAs from ovaries (Fig. 1A), RT-PCR, quantitative PCR, and *in situ* hybridization analyses showed that the short version of *Mad2* mRNA was dominant in oocytes (Figs. 1A-B and S1A). FISH analysis showed that *Mad2* mRNA was distributed in the oocyte cytoplasm by forming RNA granules (Fig. 1C). The amount of Mad2 as well as that of Cyclin B1 increased after resumption of meiosis (Fig. 1D). Consistent with this, poly(A) tails of *Mad2* mRNA were elongated 4 h after resumption of meiosis as in the case of *Cyclin B1* (Fig. 1E). Inhibition of protein synthesis with puromycin prevented the accumulation of Mad2 in oocytes even when meiosis had resumed (Fig. S1B and C). Taken together, the results indicate that Mad2 protein is accumulated in an early period of oocyte maturation by the translational activation of dormant mRNA stored in oocytes.

**Figure 1.**
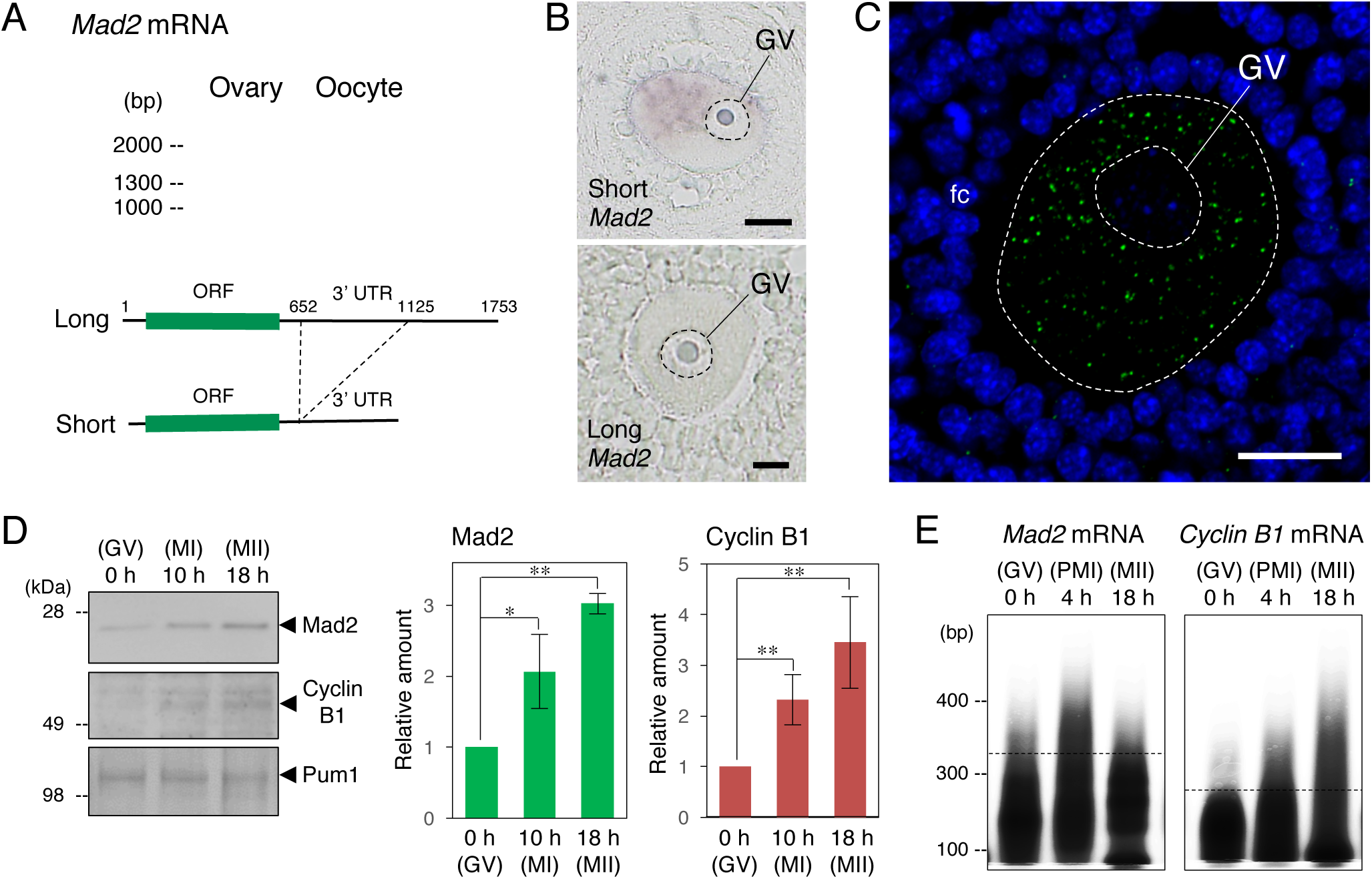
Expression and translational regulation of *Mad2* mRNA in mouse oocytes. **(A**, top) RT-PCR amplification for *Mad2* mRNA in the mouse ovary and oocyte. Similar results were obtained from three independent experiments. (bottom) Schematic views of long and short *Mad2* mRNAs. **(B)** Detection of *Mad2* mRNA in oocytes by *in situ* hybridization. Similar results were obtained from three independent experiments. **(C)** FISH analysis of *Mad2* mRNA (green). DNA is shown in blue. Similar results were obtained from three independent experiments. **(D**, left) Immunoblotting of Mad2, Cyclin B1 and Pum1 in oocytes at 0, 10, and 18 h after resumption of meiosis. (right) Quantitative analysis (mean ± SD; n = 3). *t*-test: \**P* < 0.05, \*\**P* < 0.01. **(E)** Poly(A) tail analysis of *Mad2* and *Cyclin B1* mRNAs in oocytes at 0, 4, and 18 h after resumption of meiosis. Similar results were obtained from three independent experiments. GV, germinal vesicle; fc, follicle cells. Bars: 20 *µ*m.

### *Mad2* mRNA is a Pum1-target mRNA and forms granules distinct from *Cyclin B1* RNA granules

We then assessed the mechanism by which the translation of *Mad2* mRNA is temporally regulated. Since *Mad2* mRNA was translated in a period similar to that for *Cyclin B1* mRNA and contains several putative Pumilio-binding elements (PBEs) in its 3’UTR (Fig. S2A), we investigated whether Pum1 binds to *Mad2* mRNA by using an immunoprecipitation assay followed by RT-PCR. *Mad2* and *Cyclin B1* mRNAs, but not α*-tubulin* and *β-acti*n mRNAs, were detected in precipitations with an anti-Pum1 antibody, while neither of them was detected in precipitations with control IgG (Fig. 2A), indicating that Pum1 targets *Mad2* mRNA as well as *Cyclin B1* mRNA. From these results, we speculated that both mRNAs were assembled into the same granules. However, double FISH analysis showed that the two mRNAs formed distinct granules (Fig. 2B). Notably, granules containing *Mad2* or *Cyclin B1* mRNA were found to be distributed close to each other (Fig. 2B, arrows).

**Figure 2.**
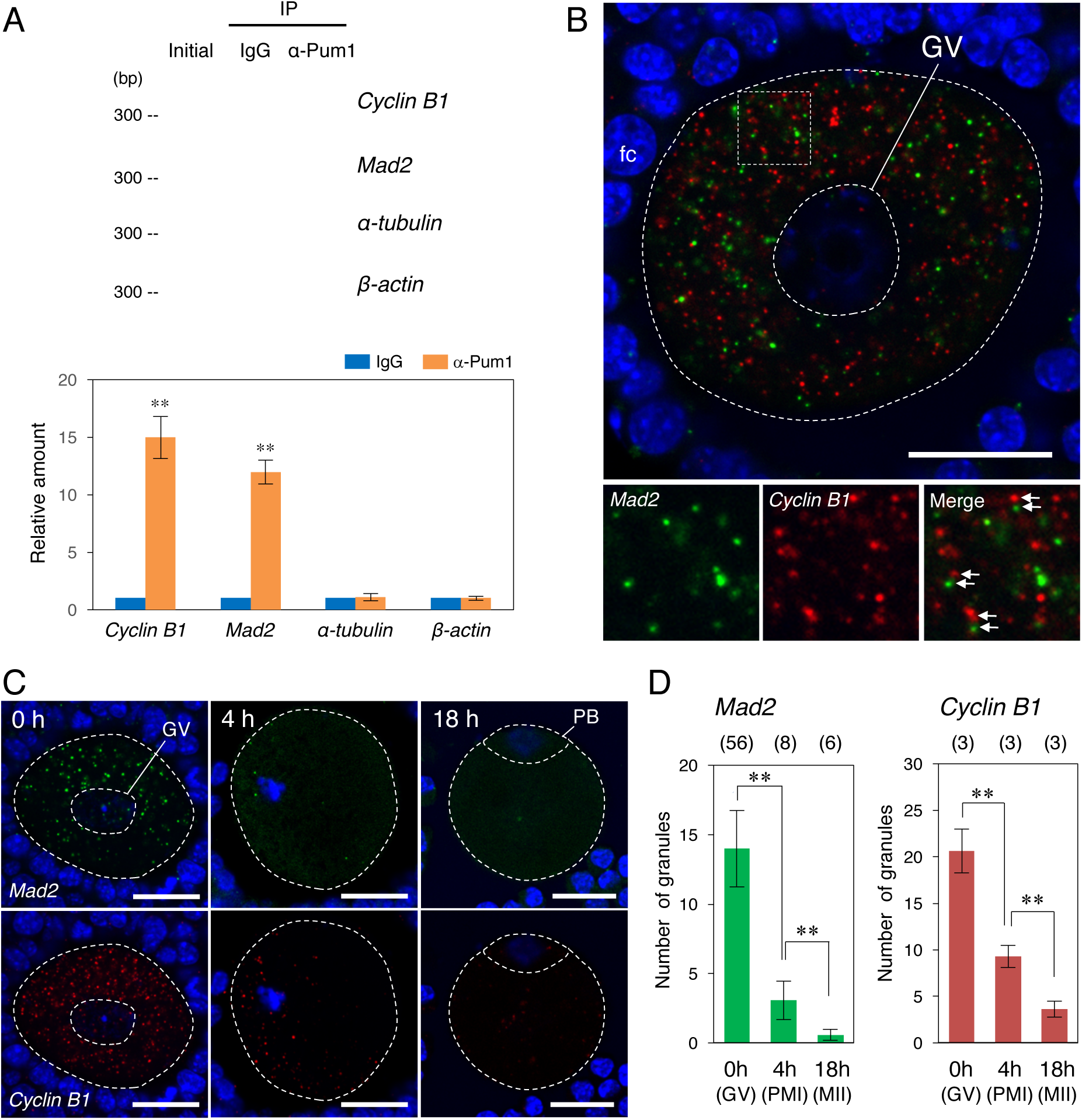
Interaction with Pum1 and cytoplasmic regulation of *Mad2* mRNA in mouse oocytes. **(A**, top) Semi-quantitative RT-PCR of ovary extracts before IP (Initial) and IP with goat IgG (IgG) or anti-Pum1 goat antibody (α-Pum1) for *Cyclin B1, Mad2*, α*-tubulin*, and *β-actin* transcripts. (bottom) Quantitative analysis (mean ± SD; n = 3). *t*-test: \*\**P* < 0.01. **(B)** FISH analysis of *Mad2* (green) and *Cyclin B1* (red) mRNAs in a mouse oocyte. DNA is shown in blue. (insets) Enlarged views of the boxed region. Arrows indicate *Mad2* and *Cyclin B1* RNA granules distributed closely to each other. Similar results were obtained from three independent experiments. **(C)** FISH analysis of oocytes at 0, 4, and 18 h after resumption of meiosis. **(D)** The numbers of RNA granules per 100 *µ*m^2^ in individual oocytes at 0, 4, and 18 h were counted (mean ± SD). The numbers in parentheses indicate the total numbers of oocytes analyzed. *t*-test: \*\**P* < 0.01. GV, germinal vesicle; fc, follicle cells; PB, polar body. Bars: 20 *µ*m.

Time course analysis showed that the number of *Mad2* RNA granules was decreased at 4 h (prometaphase I) and that the granules had almost completely disappeared at 18 h (metaphase II) after resumption of meiosis, being consistent with the changes in *Cyclin B1* RNA granules (Fig. 2C and D) (Kotani et al., 2013). These results suggest that translation of *Mad2* mRNA is temporally regulated through formation and disassembly of RNA granules, similar to the cytoplasmic regulation of *Cyclin B1* mRNA (Kotani et al., 2013).

### Pum1 forms aggregates that surround target mRNAs

To further assess the mechanism by which translation of *Mad2* and *Cyclin B1* mRNAs is temporally regulated by Pum1, we analyzed the distribution of Pum1 in the oocyte cytoplasm. Immunofluorescence analysis showed that Pum1 was ununiformly distributed in the cytoplasm of immature oocytes and appeared to form aggregates in highly clustered structures (Fig. 3A). FISH analysis showed that Pum1 aggregates surrounded and partially overlapped *Cyclin B1* and *Mad2* RNA granules (Fig. 3B). To assess the molecular mechanisms of Pum1 aggregation, we then examined the distribution of GFP-Pum1 and mutant forms of Pum1 by injecting mRNA into mouse oocytes. GFP-Pum1 was distributed in a way similar to that of endogenous Pum1, i.e., it appeared to form highly clustered aggregates (Fig. 3C and D) surrounding *Cyclin B1* and *Mad2* RNA granules (Fig. S2B). Pum1 contains a glutamine/asparagine (Q/N)-rich domain (Fig. 3E), also identified as a prion-like domain (Fig. S2C; Lancaster et al., 2014), which is thought to promote highly ordered aggregation of proteins (Lancaster et al., 2014; Salazar et al., 2010). GFP-Pum1 that lacks the Q/N-rich domain (GFP-Pum1ΔQN) (Fig. 3E) was distributed uniformly throughout the oocyte cytoplasm (Fig. 3C). Taken together, the results indicate that Pum1 assembles into aggregates by the Q/N-rich domain, and these aggregates seem to cover target mRNAs.

**Figure 3.**
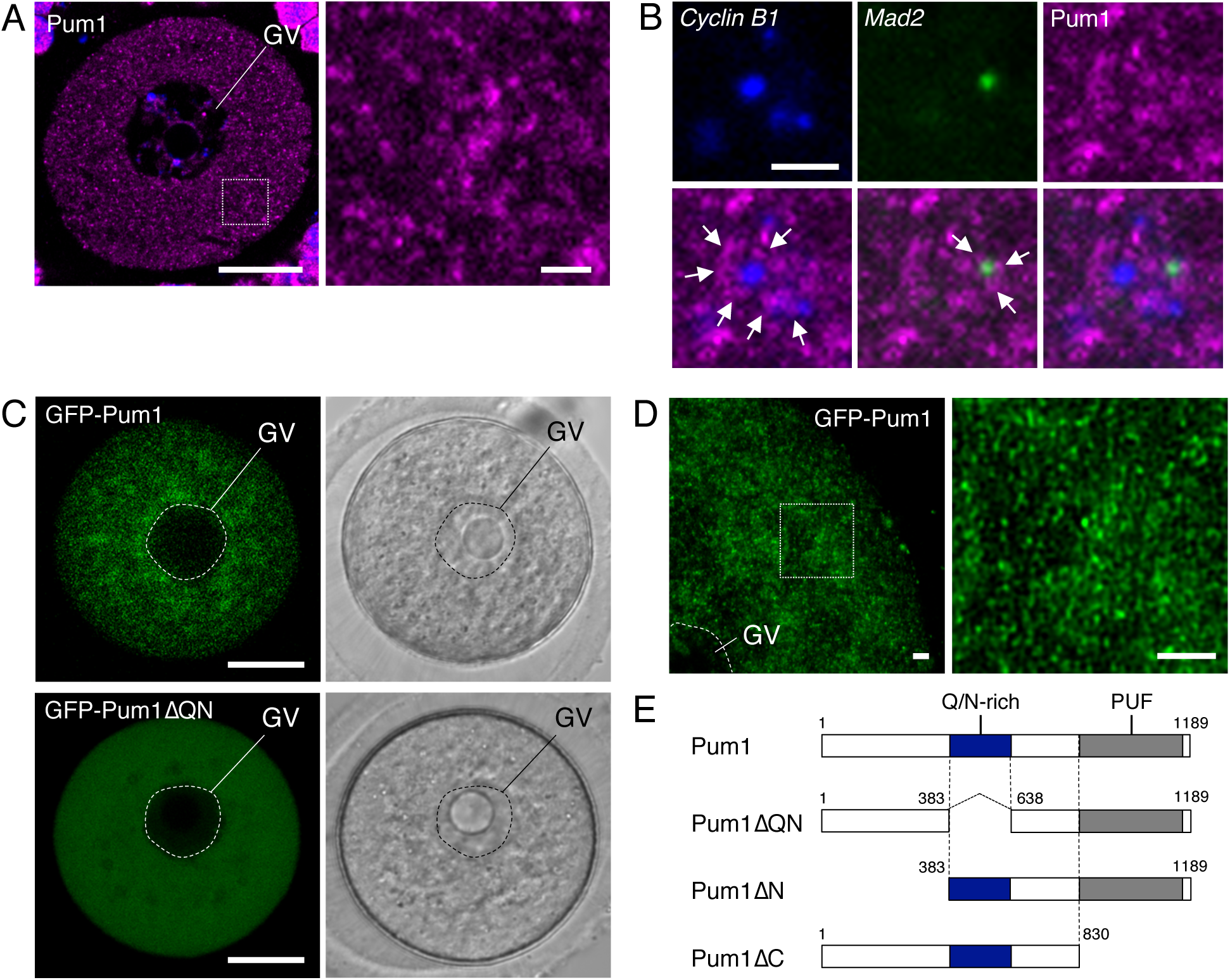
Formation of Pum1 aggregates that surround *Cyclin B1* and *Mad2* RNA granules. **(A**, left) Immunofluorescence of Pum1 in immature oocytes. DNA is shown in blue. (right) An enlarged view of the boxed region. Similar results were obtained from three independent experiments. **(B)** FISH analysis of *Cyclin B1* (blue) and *Mad2* (green) mRNAs and immunostaining of Pum1 (magenta) in immature oocytes. Arrows indicate Pum1 aggregates surrounding *Cyclin B1* and *Mad2* RNA granules. Similar results were obtained from three independent experiments. **(C)** Distributions of GFP-Pum1 (top) and GFP-Pum1ΔQN (bottom). Images in a bright field are shown on the right. **(D**, left) A high-resolution image of GFP-Pum1. (right) An enlarged view of the boxed region. Similar results were obtained from six independent experiments. **(E)** Schematic diagrams of Pum1, Pum1ΔQN, Pum1ΔN, and Pum1ΔC. GV, germinal vesicle. Bars: 20 *µ*m in A (left) and C, 2 *µ*m in A (right), B and D.

We then analyzed the distribution of Pum1 lacking the N-terminus (GFP-Pum1ΔN) or lacking the C-terminus, which contains the PUF domain responsible for binding to target mRNAs (Zhang et al., 1997) (GFP-Pum1ΔC: Fig. 2E). GFP-Pum1ΔN formed aggregates similar to those of GFP-Pum1 (Fig. S2D and Fig. 6A). In contrast, GFP-Pum1ΔC formed aggregates larger than those of GFP-Pum1 (Fig. S2D and Fig. 6A), indicating that the C-terminus PUF domain is involved in regulating the size of aggregates.

**Figure 4.**
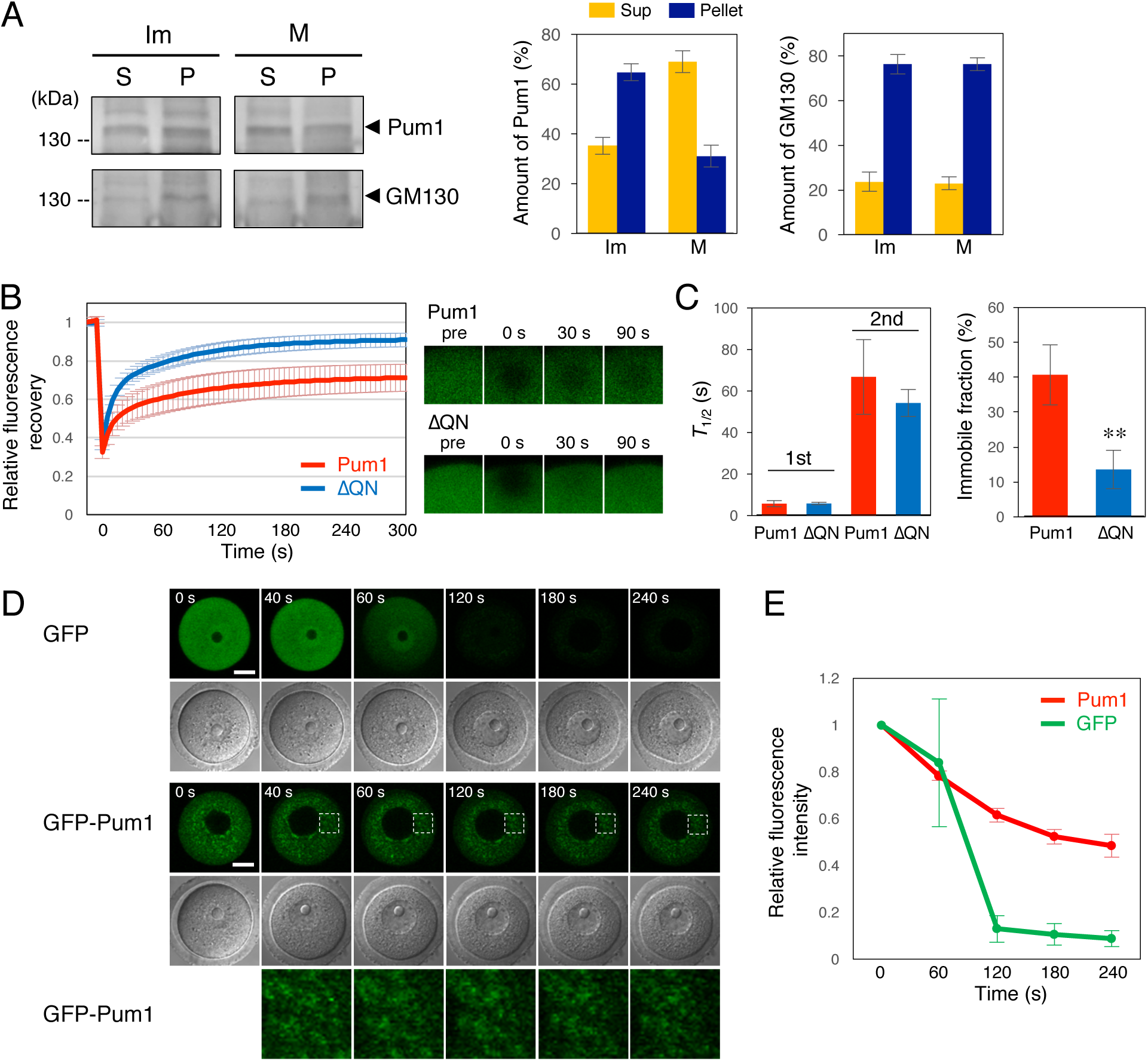
Solid-like properties of Pum1 in immature oocytes. **(A**, left) Ultracentrifugation analysis of Pum1. Immature (Im) and mature (M) zebrafish oocytes were centrifuged, and the supernatant (S) and pellet (P) equivalent to one oocyte were analyzed by immunoblotting. GM130 is a Golgi matrix protein. (right) Quantitative analysis of Pum1 and GM130 (means ± SD; n = 3). **(B)** FRAP analysis of GFP-Pum1 (Pum1) and GFP-Pum1ΔQN (ΔQN) in immature mouse oocytes. Fluorescence recovery curves for GFP-Pum1 (n = 12) and GFP-Pum1ΔQN (n = 14) are shown (mean ± SD). **(C)** Values of *t*_*1/2*_ (left) and percentages of immobile fractions of GFP-Pum1 and GFP-Pum1ΔQN (right). *t*-test: \*\**P* < 0.01. (**D**) Time course of GFP and GFP-Pum1 after permeabilization with digitonin. Similar results were obtained in 11 oocytes from two independent experiments. Bars: 20 *µ*m. (**E**) Quantitative analysis of fluorescence intensity in D (mean ± SD; n = 3).

**Figure 5.**
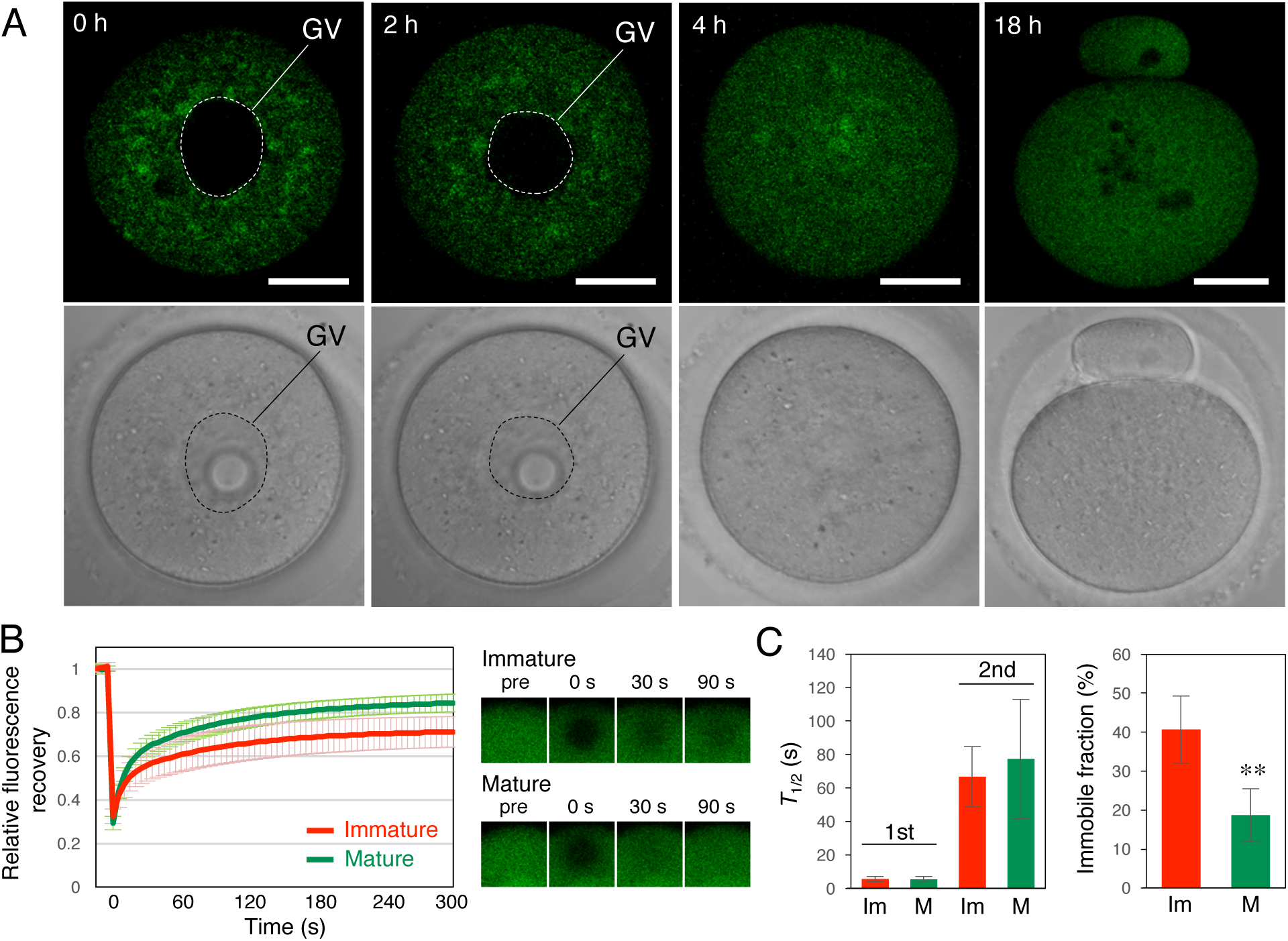
Solid-like properties of Pum1 are changed during oocyte maturation. **(A)** Time course of GFP-Pum1 at 0, 2, 4 and 18 h after resumption of meiosis. Similar results were obtained from six independent experiments. GV, germinal vesicle. Bars: 20 *µ*m. **(B)** FRAP analysis of GFP-Pum1 in immature and mature mouse oocytes. Fluorescence recovery curves in immature (n = 12) and mature (n = 6) oocytes are shown (mean ± SD). **(C)** Values of half time of recovery (*t*_*half*_) (left) and percentages of immobile fractions of GFP-Pum1 (right) in immature (Im) and mature (M) oocytes. *t*-test: \*\**P* < 0.01.

**Figure 6.**
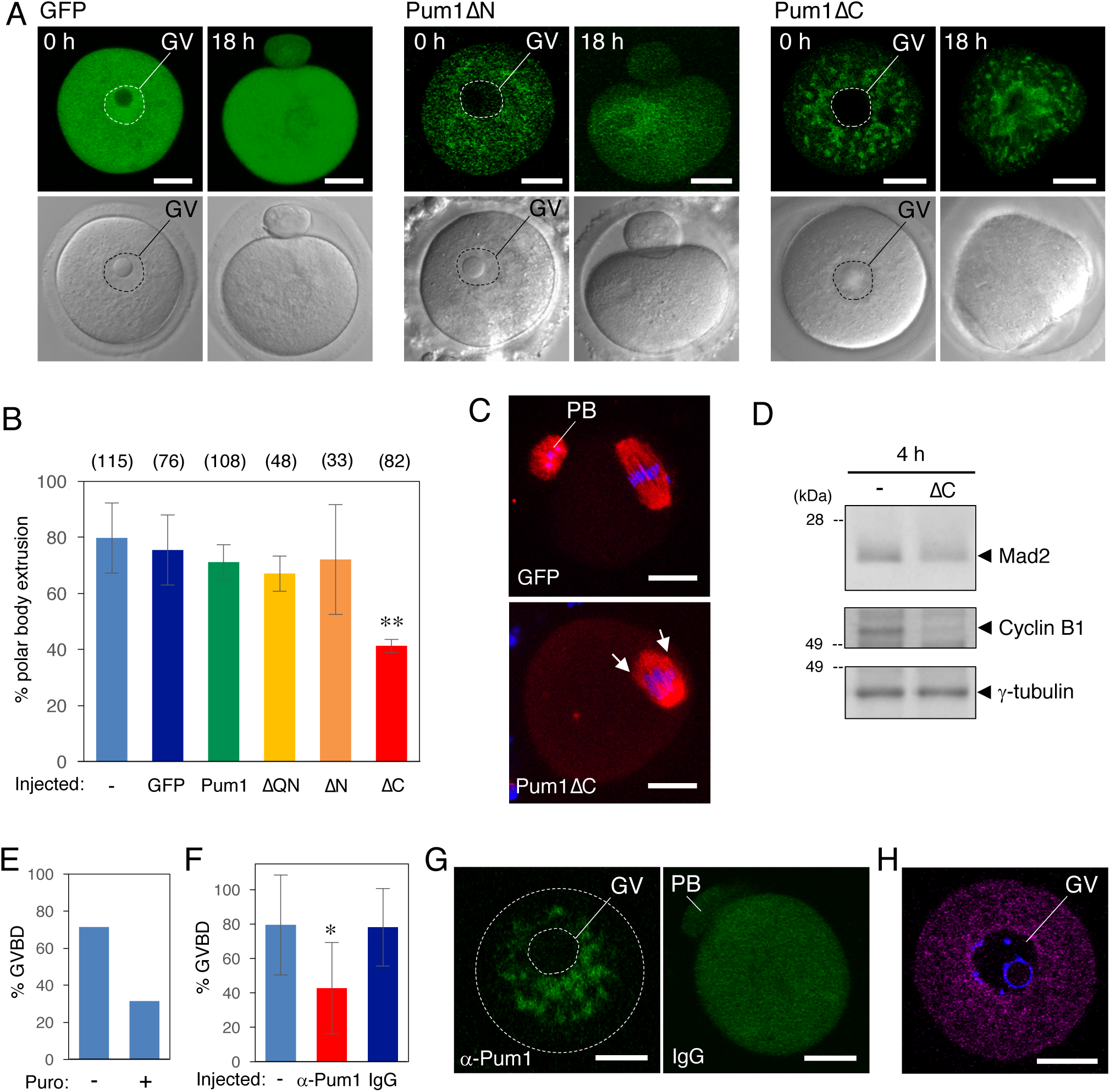
Stabilization of Pum1 aggregates prevents the translation of target mRNA. **(A)** Distributions of GFP, GFP-Pum1ΔN and GFP-Pum1ΔC at 0 and 18 h after resumption of meiosis. **(B)** Percentages of oocytes not injected (-) and injected with GFP, GFP-Pum1 (Pum1), GFP-Pum1ΔQN (ΔQN), GFP-Pum1ΔN (ΔN), and GFP-Pum1ΔC (ΔC) that extruded a polar body (means ± SD; n = 3). The numbers in parentheses indicate the total numbers of oocytes analyzed. *t*-test relative to the oocytes injected with GFP: \*\**P* < 0.01. **(C)** Immunofluorescence of *β*-tubulin (red) in oocytes injected with GFP or GFP-Pum1ΔC (Pum1ΔC). DNA is shown in blue. Arrows indicate multiple poles. Similar results were obtained from three independent experiments. **(D)** Immunoblotting of Mad2, Cyclin B1 and γ-tubulin in oocytes not injected (-) and injected with GFP-Pum1ΔC (ΔC) 4 h after resumption of meiosis. Similar results were obtained from three independent experiments. **(E)** Percentages of oocytes incubated with (+) and without (-) puromycin (Puro) that induced GVBD. **(F)** Percentages of oocytes not injected (-) and injected with anti-Pum1 antibody (α-Pum1) or control IgG (IgG) that induced GVBD (means ± SD; n = 5). *t*-test: \**P* < 0.05. **(G)** Distribution of GFP-Pum1 in oocytes injected with anti-Pum1 antibody (α-Pum1) or control IgG (IgG). (**H**) Distribution of the injected anti-Pum1 antibody (magenta). DNA is shown in blue. GV, germinal vesicle; PB, polar body. Bars: 20 *µ*m.

### Pum1 shows insoluble and immobile properties in immature oocytes

We then examined the properties of endogenous Pum1 by ultracentrifugation. Since we were unable to obtain appropriate amounts of materials by using mouse oocytes, we used zebrafish oocytes for this analysis. Zebrafish Pum1 has been shown to target *cyclin B1* mRNA (Kotani et al., 2013) and it contains the Q/N-rich domain also identified as a prion-like domain (Fig. S2C). Ultracentrifugation analysis showed that most of the endogenous Pum1 (64.8% ± 3.4%, n = 3) was concentrated in an insoluble fraction in immature oocytes (Fig. 4A), supporting the results of immunofluorescence showing that endogenous Pum1 forms aggregates (Fig. 3). These results suggest that highly clustered Pum1 aggregates exhibit a solid-like property.

We next examined the properties of GFP-Pum1 in mouse oocytes by FRAP analysis. As a control, GFP-Pum1ΔQN was analyzed. After photobleaching, the fluorescence of GFP-Pum1 and GFP-Pum1ΔQN gradually recovered (Fig. 4B). The fluorescence recovery curves were fitted to a double exponential association model. The half time of recovery (*t*_1/2_) of the first fraction of GFP-Pum1 was rapid, while that of the second fraction of GFP-Pum1 was slow (Fig. 4C, left), suggesting that a part of Pum1 forms large complexes. Moreover, a critical finding was that a significant fraction of GFP-Pum1 (40.7% ± 8.6%, n = 12) showed immobility (not recovering after photobleaching), while only a small fraction of GFP-Pum1ΔQN (13.6% ± 5.5%, n = 14) was static (Fig. 4B and C, right). Thereby, the Q/N-rich region promotes the assembly of Pum1 into highly ordered aggregates in an immobile state.

We further analyzed the properties of Pum1 by permeabilizing oocytes with digitonin. GFP rapidly diffused out of the oocytes after permeabilization (Fig. 4D and E). In contrast, the structure and intensity of GFP-Pum1 aggregates persisted after permeabilization (Fig. 4D and E). Taken together, the immunofluorescence, ultracentrifugation, FRAP and permeabilization analyses demonstrate that Pum1 assembles into aggregates in a solid-like state in immature oocytes. A recent study demonstrated that GFP-Pum1 forms solid-like substructures of RNA granules in human culture cells (Shiina, 2019), being consistent with our results in oocytes.

### Pum1 aggregates are dissolved prior to translational activation of target mRNAs

We next examined whether the distribution and properties of Pum1 changed during oocyte maturation. Time course analysis of GFP-Pum1 showed that the Pum1 aggregates disappeared after resumption of meiosis (Fig. 5A). Most of the aggregates of GFP-Pum1 had disappeared 4 h after resumption of meiosis, at which time poly(A) tails of *Mad2* and *Cyclin B1* mRNA were elongated (Fig. 1E) and the granules of both RNAs had disappeared (Fig. 2C), suggesting a link between translational activation of target mRNAs and Pum1 dissolution. Consistent with these observations, the ultracentrifugation assay showed that a large part of endogenous Pum1 became soluble (69.0% ± 4.4%, n = 3) in mature oocytes, compared with the soluble fraction in immature oocytes (35.2% ± 3.4%, n = 3) (Fig. 4A). FRAP analysis in mouse oocytes indicated that the *t*_1/2_ of GFP-Pum1 was not significantly different between immature and mature oocytes (Fig. 5B and C, left). In contrast, the percentage of immobile fractions of GFP-Pum1 was significantly reduced in mature oocytes (18.8% ± 6.8%, n = 6) compared with that in immature oocytes (40.7% ± 8.6%, n = 12) (Fig. 5B and C, right). Taken together, the results indicate that Pum1 aggregates dissolve during oocyte maturation and suggest that the change in the property of Pum1 from insoluble, solid-like to soluble, liquid-like is crucial for temporal regulation of target mRNA translation.

### Stabilization of Pum1 aggregates prevents the translation of target mRNAs

We next assessed whether the change in the property of Pum1 was involved in the translational regulation of target mRNAs. By observing the distributions of truncated forms of Pum1 after resumption of meiosis, we found that the large aggregates of GFP-Pum1ΔC were stable and persisted until 18 h (Fig. 6A). In contrast, GFP-Pum1ΔQN no longer formed aggregates (Fig. S3A), and the aggregates of GFP-Pum1ΔN were dissociated within 4 h (Fig. S3B and Fig. 6A). Consistent with the observations after resumption of meiosis, GFP-Pum1, Pum1ΔQN, and Pum1ΔN did not affect the progression of oocyte maturation, while GFP-Pum1ΔC prevented polar body extrusion (Fig. 6A and B). Temporal synthesis of proteins is required for proper spindle formation in meiosis I (Davydenko et al., 2013; Kotani and Yamashita, 2002; Polanski et al., 1998; Susor et al., 2015). In oocytes expressing GFP-Pum1ΔC, meiosis I spindles were defective and synthesis of Mad2 and Cyclin B1 was attenuated (Fig. 6C and D). These results suggest that insoluble GFP-Pum1ΔC inhibited translational activation of Pum1-target mRNAs by stabilizing Pum1 aggregates, resulting in failure in spindle formation and polar body extrusion. Since Pum1 targets thousands of mRNAs in the testis and brain (Chen et al., 2012; Zhang et al., 2017), syntheses of many proteins responsible for correct spindle formation would be attenuated in oocytes expressing GFP-Pum1ΔC.

We then examined the effects of Pum1 inhibition on the progression of oocyte maturation by injecting the anti-Pum1 antibody. To effectively analyze the effect of the anti-Pum1 antibody, we incubated oocytes with 1 *µ*M milrinone, which partially prevents resumption of meiosis. Under this condition, 50-90% of the oocytes underwent germinal vesicle breakdown (GVBD) (Fig. 6E and F) in a manner dependent on protein synthesis (Fig. 6E). Injection of the anti-Pum1 antibody, but not control IgG, prevented GVBD and dissolution of GFP-Pum1 aggregates (Fig. 6F and G). The injected anti-Pum1 antibody was distributed within the cytoplasm in a way similar to that of endogenous Pum1 (Fig. 6H). These results strongly suggest that the anti-Pum1 antibody inhibited the dissolution of endogenous Pum1 aggregates and thereby prevented the translational activation of Pum1-target mRNAs.

### Pum1 phosphorylation promotes the dissolution of aggregates

We finally assessed the mechanism by which Pum1 aggregates are dissolved. As observed in *Xenopus* and zebrafish (Ota et al., 2011; Saitoh et al., 2018), the electrophoretic mobility of Pum1 was reduced in mature mouse oocytes (Fig. 7A, left). This reduction was recovered by phosphatase treatment (Fig. 7A, right), indicating that Pum1 is phosphorylated during mouse oocyte maturation. Treatment of immature oocytes with okadaic acid (OA), a protein phosphatase 1 and 2A (PP1 and PP2A) inhibitor, induced Pum1 phosphorylation and rapid dissolution of Pum1 aggregates (Fig. 7B-D). These results suggest that kinases responsible for Pum1 phosphorylation are present and at least partially active in immature oocytes. Polo-like kinase (Plk) 1 and 4 were shown to be present in immature mouse oocytes (Bury et al., 2017; Pahlavan et al., 2000). Interestingly, inhibition of Plk4, but not that of Plk1, prevented the dissolution of Pum1 aggregates (Figs. 7C-D and S3C). Inhibition of Plk4 also prevented the phosphorylation of Pum1 (Fig. 7E). These results indicate that Plk4-mediated phosphorylation of Pum1 promotes dissolution of Pum1 aggregates.

**Figure 7.**
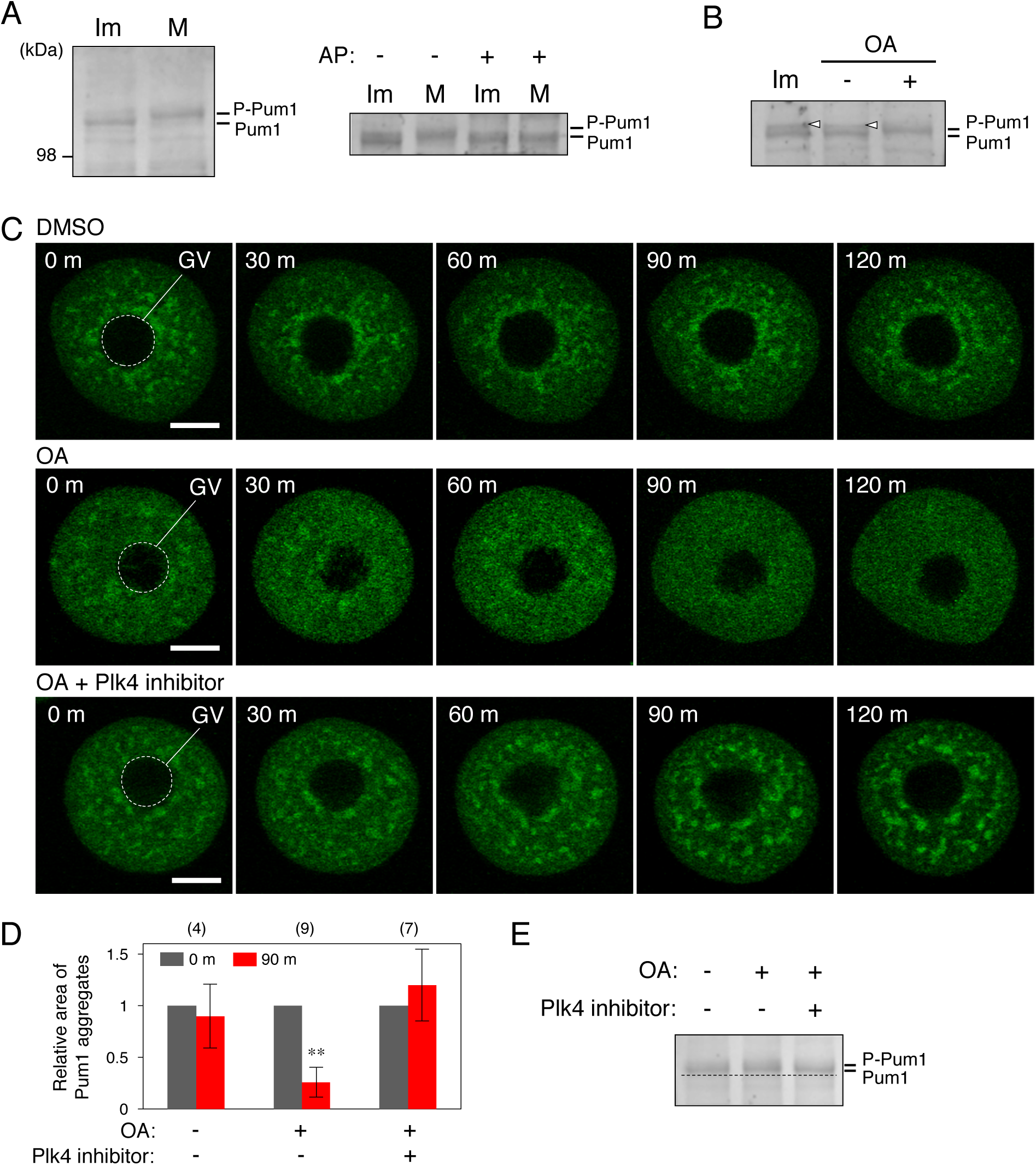
Phosphorylation of Pum1 triggers the dissolution of aggregates. **(A)** Phosphorylation of Pum1 (P-Pum1). (left) Immature (Im) and mature (M) oocytes were analyzed by immunoblotting. (right) Treatment with (+) and without (-) alkaline phosphatase (AP). Similar results were obtained from two independent experiments. **(B)** Pum1 phosphorylation in oocytes treated with OA (+) or DMSO (-). Arrowheads show nonspecific bands. Similar results were obtained from three independent experiments. **(C)** Time course of GFP-Pum1 in oocytes treated with DMSO, OA, or OA and Plk4 inhibitor 0-120 min after treatment. Similar results were obtained from three independent experiments. (**D**) Quantitative analysis of Pum1 aggregates in oocytes treated with (+) and without (-) OA or Plk4 inhibitor. The numbers in parentheses indicate the total numbers of oocytes analyzed. *t*-test: \*\**P* < 0.01. **(E)** Pum1 phosphorylation in oocytes at 60 min after treatment with (+) and without (-) OA or Plk4 inhibitor. The dotted line indicates the basal size of Pum1. GV, germinal vesicle. Bars: 20 *µ*m.

## Discussion

Extensive biochemical studies have demonstrated the importance of *cis*-acting mRNA elements and *trans*-acting RNA-binding proteins in the regulation of temporal translation (Radford et al., 2008). However, their cytoplasmic and molecular mechanisms remain largely unknown. Our results provide an aggregation-dissolution model for temporal and spatial control of mRNA translation, i.e., Pum1 aggregates in clustered solid-like structures ensure translational repression of target mRNAs by stably maintaining their granular structures, and the dissolution of aggregates into a liquid-like state by phosphorylation permits the disassembly of granules and translational activation of mRNAs. Given that many dormant mRNAs stored in oocytes contain PBEs (Chen et al., 2011) and Pum1 targets more than one thousand mRNAs in the testis and brain (Chen et al., 2012), Pum1 would target a large number of mRNAs in oocytes. In addition, Pum1 aggregates might be comprised of these target mRNAs and related proteins and thereby allow their coordinated regulation. Our results will be a basis for understanding how translational timings of hundreds of mRNAs are coordinately regulated.

### Phase changes of Pum1 and translational regulation of target mRNAs

Recent studies have demonstrated that many of the RNA-binding proteins harbor prion-like domains and that some of these proteins have the ability to assemble RNA granules (Decker et al., 2007; Gilks et al., 2004; Reijns et al., 2008). These RNA-binding proteins were shown to promote liquid-liquid phase separation, resulting in the assembly of protein-RNA complexes into droplets (Elbaum-Garfinkle et al., 2015; Lin et al., 2015; Molliex et al., 2015; Nott et al., 2015). These droplets are thought to function as partitions that effectively maintain stability and/or translational repression of mRNAs. In contrast, phase transition of the liquid droplets into solid-like structures such as amyloid fibrils has been thought to contribute to pathological diseases such as amyotrophic lateral sclerosis (ALS) (Li et al., 2013; Weber and Brangwynne, 2012). However, more recently, solid granules were found to assemble during muscle regeneration in a physical state (Vogler et al., 2018). In addition, core regions of stress granules were shown to exhibit solid-like properties (Jain et al., 2016; Shiina, 2019; Wheeler et al., 2016). Although these findings suggest the involvement of solid granules in RNA regulation, the physiological importance of the phase changes of protein aggregation from liquid to solid states and vice versa remains unclear.

In this study, we demonstrated that Pum1 assembled into aggregates in highly clustered structures through the Q/N-rich region and these aggregates showed solid-like properties in immature oocytes (Figs. 3 and 4). After initiation of oocyte maturation, the Pum1 aggregates dissolved into a liquid-like state (Figs. 4A and 5). The mutant form of Pum1 that lacks the C-terminal PUF domain, Pum1ΔC, is expected to be unable to bind to target mRNAs but to have the ability to form assemblies via the Q/N-rich region. Since an RNA molecule was shown to buffer the assembly of RNA-binding proteins that harbor prion-like domains into a solid-like aggregates (Maharana et al., 2018), it is possible that the lack of RNA-binding ability of Pum1ΔC resulted in the assembly of large and stable aggregates (Figs. S2D and 6A). Pum1ΔC would stabilize endogenous Pum1 aggregates via the Q/N-rich region-mediated aggregation and thereby prevent the translational activation of Pum1-target mRNAs (Fig. 6A-D). The anti-Pum1 antibody also prevented dissociation of Pum1 aggregates (Fig. 6E-H). One possible explanation for this is that the binding of Pum1 antibodies attenuated the phosphorylation of Pum1. Another possibility is that the antibody affected the conformation or composition of Pum1 assemblies, preventing aggregate dissolution and translational activation of mRNAs. Collectively, our results demonstrated a physiological significance of phase changes of protein aggregation in translational repression and activation of target mRNAs.

### Regulation of the subcellular structures and states of Pum1 by phosphorylation and dephosphorylation

P granules are the germinal granules in *C. elegance* that are important for fate decision of germline cells. Live imaging of embryos demonstrated that P granules behave as dynamic liquid droplets (Brangwynne et al., 2009). Intriguingly, disassembly of P granules after fertilization was shown to require MBK-2 kinase, while subsequent assembly of P granules at the posterior region of embryos required protein phosphatase 2A (PP2A) (Gallo et al., 2010; Wang et al., 2014). MEG-1 and MEG-3 were found to be the substrates of MBK-2 and PP2A in the granules (Wang et al., 2014). These results demonstrated that the dynamics of liquid RNA granules is regulated by phosphorylation and dephosphorylation of assembled proteins.

Our results suggest the importance of protein phosphorylation and dephosphorylation for changes in structures and states of solid-like aggregates. SDS-PAGE analysis demonstrated that Pum1 was phosphorylated during mouse oocyte maturation (Fig. 7A). Interestingly, treatment of oocytes with OA, an inhibitor of PP1 and PP2A, rapidly dissociated Pum1 aggregates and induced Pum1 phosphorylation (Fig. 7B-D). Since PP2A was shown to be localized in the cytoplasm of GV-stage mouse oocytes, while PP1 was dominantly localized in the nucleus (Smith et al., 1998), PP2A would be a phosphatase involved in Pum1 dephosphorylation and the maintenance of Pum1 aggregates. Even when the activity of PP1 and PP2A was inhibited by OA, Pum1 was not phosphorylated and the aggregates persisted in the presence of a Plk4 inhibitor (Fig. 7C-E), suggesting that Plk4 is a kinase responsible for Pum1 phosphorylation and aggregate dissolution. However, other kinases would phosphorylate Pum1, since inhibition of Plk4 activity delayed, but did not completely prevent, the disolution of Pum1 aggregates and Pum1 phosphorylation after initiation of oocyte maturation (unpublished data). Puf3, one of the PUF family proteins in yeast, was shown to be phosphorylated at up to 20 sites throughout the entire region (Lee and Tu, 2015). In addition, we previously showed that Pum1 was phosphorylated at multiple sites in an early period of oocyte maturation in zebrafish (Saitoh et al., 2018). Although the phosphorylation sites responsible for the aggregate dissolution remain to be identified, these results suggest that many sites including those in the Q/N-rich domain might be phosphorylated, resulting in Pum1 aggregate dissolution.

### Subcellular structures of Pum1 and homogenous RNA granules

An intriguing finding in this study is that Pum1-target *Mad2* and *Cyclin B1* mRNAs formed distinct granules in the oocyte cytoplasm, instead of making granules containing both mRNAs (Fig. 2). Pum1 was found to produce highly clustered structures that surrounded both *Mad2* and *Cyclin B1* RNA granules (Fig. 3). These structures partially resemble those of germ granules in *Drosophila* embryos, in which mRNAs form homogenous RNA clusters and are spatially positioned within the granules, while RNA-binding proteins are evenly distribute throughout the granules (Trcek et al., 2015). These findings suggest the existence of a common mechanism by which each mRNA could be organized into homogenous particles. However, in contrast to our findings, the structures of germ granules were not changed during early stages of embryogenesis and were independent of the control of mRNA translation and degradation (Trcek et al., 2015). Therefore, the function of spacially organized structures of germ granules in *Drosophila* embryos seems to be different from the function of subcellular structures of Pum1 and RNA granules in mouse oocytes.

Our results showed that Pum1 aggregates surrounded and overlapped *Mad2* and *Cyclin B1* RNA granules but were not localized at the center of granules (Fig. 3). Given that Pum1 was shown to bind directly to PBE in the 3’UTR of target mRNAs including *Cyclin B1* (Kotani et al., 2013; Nakahata et al., 2003; Ota et al., 2011; Pique et al., 2008), Pum1-target mRNAs may compose highly ordered structures within granules, in which the 3’ ends of mRNAs are localized at the periphery of granules as in the case of a long noncoding RNA, *Neat1*, in paraspeckle nuclear bodies (Souquere et al., 2010; West et al., 2016). Details of the molecular mechansims by which Pum1 is assembled into aggregates remain unknown. One possible model is that Pum1 binds to a target mRNA via the PUF domain and subsequently assembles into aggregates via the Q/N-rich region. Another possibility is that Pum1 contains two populations; one population binds to target mRNAs and the other functions as structual scaffolds without binding to mRNAs. In addition to the homogenous assembly of Pum1, heterogenous assembly with other RNA-binding proteins may produce aggregates. In any case, the resulting Pum1 aggregates in clustered structures would make compartments that function as regulatory units with related proteins assembled together or separately. These units enable to coordinately regulate the translation of assembled mRNAs. Since Pum1 functions in diverse systems and other RNA-binding proteins that harbor prion-like domains may function in a manner similar to that of Pum1, our results will contribute to an understanding of the nature of temporal and spatial control of translation in many cell types of diverse organisms.

## Materials and Methods

### Preparation of ovaries and oocytes

All animal experiments in this study were approved by the Committee on Animal Experimentation, Hokkaido University. Mouse ovaries were dissected from 8-week-old females in M2 medium (Sigma). Oocytes were retrieved from ovaries by puncturing the ovaries with a needle in M2 medium containing 10 *µ*M milrinone, which prevents resumption of oocyte maturation. To induce oocyte maturation, the isolated oocytes were washed three times and incubated with M2 medium without milrinone at 37°C. Alternatively, oocyte maturation was induced by injection of 5 U of hCG 48 h after injection of 5 U of pregnant mare serum gonadotropin into 3-week-old females. For RT-PCR and poly(A) test (PAT) assays, ovaries and oocytes were extracted with Trizol reagent (Invitrogen) and total RNA was used for RT-PCR and RNA ligation-coupled RT-PCR. For *in situ* hybridization analysis, mouse ovaries were fixed with 4% paraformaldehyde in PBS (137 mM NaCl, 2.7 mM KCL, 10 mM Na_2_HPO_4_, and 2 mM KH_2_PO_4_, pH 7.2) (4% PFA/PBS) overnight at 4°C. For immunoblotting analysis, 30 oocytes were washed with PBS and extracted with lithium dodecylsulfate (LDS) sample buffer (Novex) at 0, 10, and 18 h after resumption of oocyte maturation. For IP/RT-PCR analysis, mouse ovaries were homogenized with an equal volume of ice-cold extraction buffer (EB: 100 mM β-glycerophosphate, 20 mM HEPES, 15 mM MgCl_2_, 5 mM EGTA, 1 mM dithiothreitol, 100 *µ*M (*p*-amidinophenyl)methanesulfonyl fluoride, 3 *µ*g/ml leupeptin; pH 7.5) containing 1% Tween20 and 100 U/ml RNasin Plus RNase Inhibitor (Promega). After centrifugation at 15,000 g for 10 min at 4°C, the supernatant was collected and used for IP.

Zebrafish ovaries were dissected from adult females in zebrafish Ringer’s solution (116 mM NaCl, 2.9 mM KCl, 1.8 mM CaCl_2_, and 5 mM HEPES; pH 7.2). Zebrafish oocytes were manually isolated from ovaries with forceps under a dissecting microscope. Oocyte maturation was induced by treatment with 1 *µ*g/ml of 17α,20β-dihydroxy-4-pregnen-3-one, a maturation-inducing hormone in fish. For ultracentrifugation analysis, fully grown immature oocytes and oocytes 3 h after MIH stimulation (matured oocytes) were homogenized with an equal volume of ice-cold EB containing 0.2% Tween20. After ultracentrifugation at 90,000 g for 30 min at 4°C, the supernatant and precipitates were collected and used for immunoblot analysis.

### RT-PCR and quantitative PCR

Total RNA extracted from mouse ovaries or 50 immature oocytes was used for cDNA synthesis using the Super Script III First Strand Synthesis System (Invitrogen). The full length of *Mad2* mRNA was amplified with the cDNA and primer sets specific to *Mad2*, m*Mad2*-f1 (5’-GTA GTG TTC TCC GTT CGA TCT AG-3’) and m*Mad2*-r1 (5’-GTA TCA CTG ACT TTT AAA GCT TGA TTT TTA-3’). The amounts of short and long *Mad2* mRNAs were quantified by using a real-time PCR system with SYBR green PCR Master Mix (Applied Biosystems) according to the manufacturer’s instructions. The short and long *Mad2* transcripts were amplified with the cDNA and primer sets to both types of *Mad2*, m*Mad2*-f2 (5’-GAA TAG TAT GGT GGC CTA CAA-3’) and m*Mad2*-r2 (5’-TTC CCT CGT TTC AGG CAC CA-3’), and primer sets specific to long *Mad2*, m*Mad2*-f3 (5’-CTG GAC CAG GAT ATA AAG AAG CG-3’) and m*Mad2*-r3 (5’-GCT GTC CTC CCT GCC TCT CT-3’). The signals obtained with distinct primer sets were normalized by standard curves obtained with plasmid DNAs encoding the short or long *Mad2* gene.

### Section *in situ* hybridization

Section *in situ* hybridization and fluorescent *in situ* hybridization (FISH) with the tyramide signal amplification (TSA) Plus DNP system (PerkinElmer) were performed according to the procedure reported previously (Takei et al., 2018). Briefly, fixed ovaries were dehydrated, embedded in paraffin, and cut into 7-*µ*m-thick sections. Digoxigenin (DIG)-labeled antisense RNA probes for the full length of short *Mad2* and sequences specific to long *Mad2* were used for detection of *Mad2* gene transcripts. No signal was detected with sense probes. After hybridization and washing, samples were incubated with an anti-DIG-horseradish peroxidase (HRP) antibody (Roche) (1:500 dilution) for 30 min. To detect *Mad2* transcripts by alkaline phosphatase (AP) staining, reaction with tyramide-dinitrophenyl (DNP) (PerkinElmer) was performed according to the manufacturer’s instructions. The samples were then incubated with an anti-DNP-AP antibody (PerkinElmer) (1:500 dilution) for 30 min, followed by reaction with NBT and BCIP according to the manufacturer’s instructions. To detect *Mad2* transcripts by fluorescence microscopy, reaction with tyramide-Fluorescein (PerkinElmer) was performed according to the manufacturer’s instructions. To detect nuclei, samples were incubated with 10 *µ*g/ml Hoechst 33258 for 10 min. After being mounted with a Prolong Antifade Kit (Molecular probes), the samples were observed under an LSM 5 LIVE confocal microscope (Carl Zeiss) at room temperature using a Plan Apochromat 63x/1.4 NA oil differential interference contrast lens and LSM 5 DUO 4.2 software (Carl Zeiss).

Double *in situ* hybridization of *Mad2* and *Cyclin B1* transcripts was performed as follows. A fluorescein-labeled antisense RNA probe for *Cyclin B1* was used for detection of the *Cyclin B1* gene transcript. Seven-*µ*m-thick sections of mouse ovaries were hybridized with a mixture of *Mad2* and *Cyclin B1* antisense RNA probes. Then the samples were incubated with an anti-Fluorescein-HRP antibody (Roche) (1:200 dilution) for 30 min. Reaction with tyramide-Cy3 (PerkinElmer) was performed according to the manufacturer’s instructions. For inactivating HRP, samples were incubated with 1% H_2_O_2_ in PBS for 15 min. Detection of the DIG-labeled antisense *Mad2* RNA probe was performed as described above. After staining with Hoechst 33258, the samples were mounted and observed under the LSM5LIVE confocal microscope. The number of *Mad2* and *Cyclin B1* RNA granules was quantified using ImageJ software, which enables detection of granules according to size (larger than 0.2 *µ*m) and intensity at the center of granules. Similar results were obtained using a fluorescein-labeled antisense RNA probe for *Mad2* and a DIG-labeled RNA probe for *Cyclin B1*.

### Immunoblotting

Mouse oocyte extracts were separated by SDS-PAGE with Bolt Bis-Tris Plus Gels (Novex), blotted onto an Immobilon membrane using a Bolt Mini Blot Module (Novex), and probed with an anti-human Pum1 goat antibody (1:1000 dillution) (Bethyl Laboratories, Inc.), an anti-human Cyclin B1 rabbit antibody (1:100 dillution) (Santa Cruz Biotechnology, Inc.), an anti-hamster Cyclin B1 mouse monoclonal antibody (1:1000 dilution) (V152, Abcam), and an anti-human Mad2 rabbit antibody (1:1000 dillution) (Bethyl Laboratories, Inc.). The supernatant and precipitates of zebrafish oocyte extracts were separated by SDS-PAGE, blotted onto an Immobilon membrane, and probed with an anti-*Xenopus* Pum1 mouse monoclonal antibody (1:1000 dillution) (Pum2A5) and an anti-GM130 mouse monoclonal antibody (1:250 dillution) (BD Biosciences). The intensity of signals was quantified using ImageJ software.

### Poly(A) test (PAT) assay

RNA ligation-coupled RT-PCR was performed according to the procedure reported previously (Kotani et al., 2013). Four hundred ng of total RNA extracted from pools of 250 mouse oocytes was ligated to 400 ng of P1 anchor primer (5’-P-GGT CAC CTT GAT CTG AAG C-NH_2_-3’) in a 10-*µ*l reaction using T4 RNA ligase (New England Biolabs) for 30 min at 37°C. The ligase was inactivated for 5 min at 92°C. Eight *µ*l of the RNA ligation reaction was used in a 20-*µ*l reverse transcription reaction using the Superscript III First Strand Synthesis System with a P1’ primer (5’-GCT TCA GAT CAA GGT GAC CTT TTT TTT-3’). Two *µ*l of the cDNA was used for the 1st PCR with the P1’ primer and an m*Mad2*-f4 primer (5’-GAC CCC ATA TTG AAA TAC ATG C-3’) or m*Cyclin B1*-f1 primer (5’-CCA CTC CTG TCT TGT AAT GC-3’) for 45 cycles. One *µ*l of the 1st PCR reaction was used for the 2nd PCR with the IRD800-P1’ primer (5’-IRD800-GCT TCA GAT CAA GGT GAC CTT TTT TTT-3’) and an m*Mad2*-f5 primer (5’-GAG CTC ACA ACG CAG TTG-3’) or m*Cyclin B1*-f2 primer (5’-CCT GGA AAA GAA TCC TGT CTC-3’) for 20 cycles. The PCR product was resolved on a 3% TAE gel and observed by using Odessay (M&S TechnoSystem). We confirmed that the increase in PCR product length was due to elongation of the poly(A) tails by cloning the 2nd PCR products and sequencing them.

### RT-PCR analysis after IP (IP/RT-PCR)

Eighty *µ*l of mouse ovary extracts was incubated with 4 *µ*l of 1.0 *µ*g/ml anti-human Pum1 goat antibody or 4 *µ*l of 1.0 *µ*g/ml control goat IgG for 1 h at 4°C. The extracts were then incubated with protein A-Sepharose beads (GE Healthcare) for 3 h at 4°C and washed five times with EB containing 1% Tween 20. After extraction of mRNAs from the beads with Trizol reagent, RT-PCR was performed using primer sets specific to *Mad2*, m*Mad2*-f6 (5’-GTG ACC ATT GTT AAA GGA ATC CAT CCC-3’) and m*Mad2*-r1, to *Cyclin B1*, m*Cyclin B1*-f3 (5’-AGT CCC TCA CCC TCC CAA AAG C-3’) and m*Cyclin B1*-r1 (5’-AAA GCT TTC CAC CAA TAA ATT TTA TTC AAC-3’), to *β-actin*, m*β-actin* -f1 (5’-AGT CCC TCA CCC TCC CAA AAG C-3’) and m*β-actin* -r1 (5’-GGT CTC AAG TCA GTG TAC AGG C-3’), and to α*-tubulin*, mα*-tubulin*-f1 (5’-CTT TGT GCA CTG GTA TGT GGG T-3’) and mα*-tubulin*-r1 (5’-ATA AGT GAA ATG GGC AGC TTG GGT-3’). The intensity of signals was quantified using ImageJ software.

### Immunofluorescence

Fixed ovaries were dehydrated, embedded in paraffin, and cut into 7-*µ*m-thick sections. After rehydration, samples were microwaved for 10 min with 0.01 M citric acid (pH 6.0) containing 0.05% Tween20, followed by cooling down for 40 min. After incubation with a TNB blocking solution (PerkinElmer) for 1 h at room temperature, the samples were incubated with anti-human Pum1 goat antibody (1:100 dilution) (Novus Biologicals) at 4°C for overnight. The samples were then incubated with anti-goat IgG-Alexa Fluor Plus 647 antibody (1:200 dilution) (Invitrogen) at room temperature for 1 h. After staining with Hoechst 33258, the samples were mounted and observed under the LSM 5 LIVE confocal microscope. No signal was detected in the reaction without the anti-Pum1 antibody. To simultaneously detect Pum1 and *Cyclin B1* and *Mad2* mRNAs, the samples were immunostained with the Pum1 antibody as described above after detection of the *Cyclin B1* and *Mad2* RNA probes in *in situ* hybridization analysis.

### mRNA injection and immunostaining

Sequences encoding the full length and parts of mouse Pum1 (ΔQN, ΔN and ΔC) were cloned into pCS2-GFP-N to produce Pum1 fused with GFP at the N terminus of Pum1. mRNAs encoding GFP, GFP-Pum1, GFP-Pum1ΔQN, GFP-Pum1ΔN, and GFP-Pum1ΔC were synthesized with an mMESSAGE mMACHINE SP6 kit (Life Technologies) and dissolved in distilled water. Ten pg of the mRNAs was injected into fully grown mouse oocytes using an IM-9B microinjector (Narishige) under a Dmi8 microscope (Leica) in M2 medium containing 10 *µ*M milrinone. After being incubated for 4 h at 37°C, the oocytes were fixed with 2% PFA/PBS containing 0.05% Triron-X100 for 1 h at 4°C for *in situ* hybridization analysis or were washed four times with M2 medium without milrinone for induction of oocyte maturation. At the appropriate time points after resumption of meiosis, the distribution of proteins fused with GFP was observed under the LSM 5 LIVE confocal microscope. To simultaneously detect GFP-Pum1 and *Cyclin B1* or *Mad2* mRNA, the fixed oocytes were attached on slide glasses using Smear Gell (GenoStaff). The oocytes were immunostained with anti-GFP mouse antibody (1:200 dilution; Roche) followed by anti-mouse IgG-Alexa Fluor 488 antibody (1:200 dilution; Molecular Probes) after hybridization and washing of the *Cyclin B1* or *Mad2* RNA probe in *in situ* hybridization analysis.

To analyze the effects of permeabilization on GFP-Pum1 aggregates, the oocytes injected with mRNA encoding GFP or GFP-Pum1 were incubated for overnight at 37°C with M2 medium containing 10 *µ*M milrinone. After observation under the LSM 5 LIVE confocal microscope, the oocytes were transferred to M2 medium containing 0.012% digitonin and 10 *µ*M milrinone. The oocytes were then observed under the confocal microscope at the appropriate time points.

To analyze the effects of GFP-Pum1ΔC on oocyte maturation, the oocytes injected with mRNA encoding GFP or GFP-Pum1ΔC were incubated for 18 h at 37°C with M2 medium and then fixed with 4% PFA/PBS for 1 h at 37°C. The samples were permeabilized with PBS containing 0.1% Triton-X100 for 20 min, followed by incubation with a blocking/washing solution (PBS containing 0.3% BSA and 0.01% Tween20) for 1 h at room temperature. The samples were then incubated with Cy3-conjugated anti-β-tubulin antibody (1:150 dilution; Sigma) for 30 min at room temperature, washed with washing solution, and mounted with VECTASHIELD Mounting Medium with DAPI (Funakoshi). The samples were observed under the LSM 5 LIVE confocal microscope.

### FRAP analysis

FRAP measurements were performed according to the procedure reported previously (Kimura and Cook, 2001; Tsutsumi et al., 2016). A Nikon Ti-E inverted microscope equipped with a Nikon A1Rsi special imaging confocal laser scanning system (Nikon) was used for the measurements. A small area (approximately 10 *µ*m diameter circle) was positioned in a region of the oocyte cytoplasm and bleached using 100% 488 nm laser with 5 scans. Images were then collected using 1.0% laser power every 5.0 s for 5.0 min. The relative fluorescence intensity in the bleached area was normalized using the intensity in the control area measured subsequently after measurement of the bleached area. The normalized intensities were analyzed using a fitting equation for a double exponential association model. A smaller bleached area (5 *µ*m diameter circle) gave equivalent results.

### Puromycin treatment and Pum1 antibody injection

To inhibit protein synthesis, oocytes were treated with 20 mM puromycin in M2 medium and incubated at 37°C. The oocytes were collected at appropriate time points after incubation with puromycin for immunoblotting analysis. Two pg of anti-Pum1 antibody was injected into fully grown mouse oocytes using the microinjector in M2 medium containing 10 *µ*M milrinone. The oocytes were then washed three times and incubated for 18 h at 37°C with M2 medium containing 1 *µ*M milrinone. To analyze the distribution of GFP-Pum1, 10 pg of the GFP-Pum1 mRNA was co-injected with 2 pg of anti-Pum1 antibody into fully grown mouse oocytes, followed by washing and incubation of oocytes as described above. The distribution of GFP-Pum1 was observed under the LSM 5 LIVE confocal microscope.

### Phosphatase treatment

The dephosphorylation experiments were performed according to the procedure reported previously (Pahlavan et al., 2000). Briefly, samples of 30 oocytes in phosphatase buffer (New England Biolabs) containing 1% SDS, 100 *µ*M (*p*-amidinophenyl)methanesulfonyl fluoride, and 3 *µ*g/ml leupeptin were incubate with 17.5 U alkaline phosphatase (New England Biolabs) at 37°C for 1 h. The reaction was stopped by adding the equal volume of LDS sample buffer. The samples were then analyzed by immunoblotting.

### Okadaic acid, BI2536, and centrinone treatment

To inhibit activities of protein phosphatase 1 and 2A, oocytes were treated with 2.5 *µ*M okadaic acid (OA) in M2 medium containing 10 *µ*M milrinone and incubated at 37°C. OA was dissolved in DMSO as stocks and diluted in M2 medium before use. As a control, oocytes were treated with DMSO. The oocytes were collected at 16 h after incubation for immunoblotting analysis. To analyze the distribution of GFP-Pum1, fully grown mouse oocytes were injected with 10 pg of the GFP-Pum1 mRNA and incubated in M2 medium containing 10 *µ*M milrinone at 37°C for 4 h, followed by treatment with OA as described above. The distribution of GFP-Pum1 was observed under the LSM 5 LIVE confocal microscope. Activities of Plk1 and Plk4 were inhibited by treating the oocytes with 100 nM BI2536 and 5 *µ*M centrinone, respectively, according to the procedure reported previously (Bury et al., 2017).

## Acknowledgements

We thank Drs. K. Kobayashi and M. Tsutsumi for technical advice on FRAP analysis. We also thank Dr. H. Maita for advice on the detection of PCR amplification in the PAT assay. This work was supported by Grant-in-Aid for Scientific Research (16K07242 to T.K.) from the Ministry of Education, Culture, Sports, Science and Technology, Japan and was in part supported by grants from Takeda Science Foundation, Daiichi Sankyo Foundation of Life Science, Suhara Memorial Foundation, and JSPS KAKENHI Grant Number JP16H06280.

## Author contributions

Conceptualization: N. Takei, Y. Takada, S. Kawamura and T. Kotani. Investigation: N. Takei, S. Kawamura, Y. Takada and A. Saitoh. Resources: J. Bormann, W.S. Yuen and J. Carroll. Project administration: T. Kotani. Writing - original draft: T. Kotani. Writing - review and editing: J. Carroll and T. Kotani.

## Conflict of interest

The authors declare that no competing interests exist.

## Supplemental figure legends

**Figure S1.**
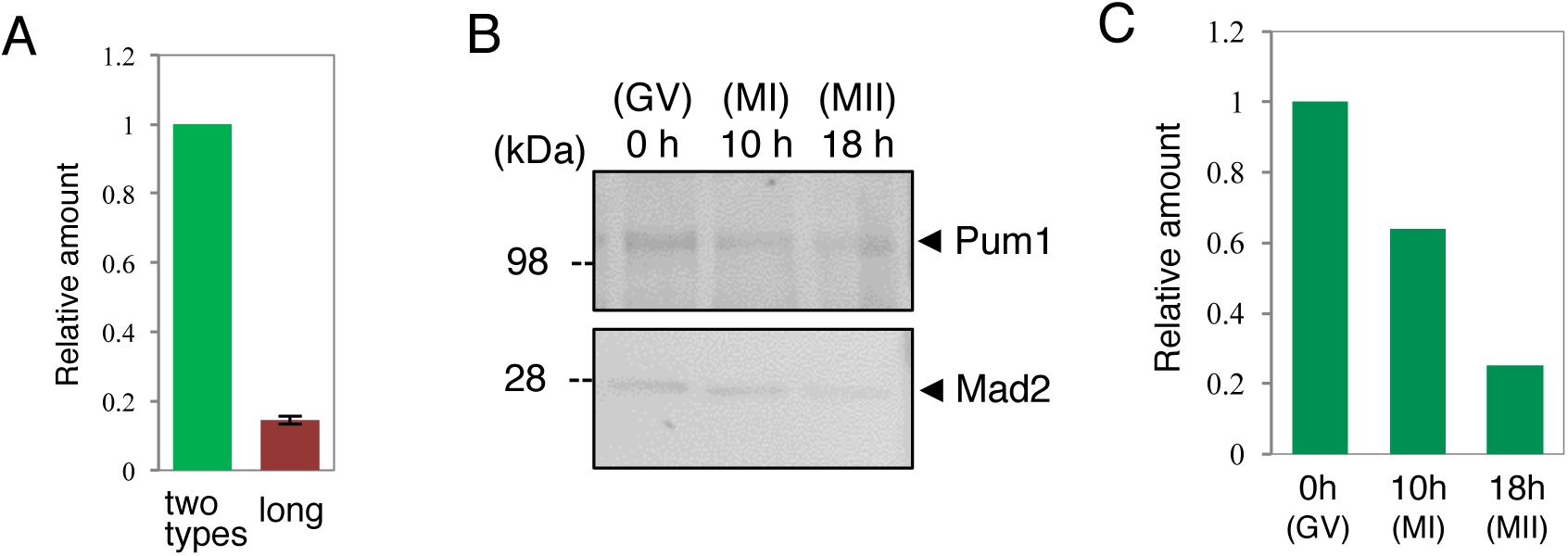
Expression of *Mad2* mRNA and effect of puromycin on Mad2 protein accumulation. **(A)** Quantitative PCR for the two types of *Mad2* mRNA and for long *Mad2* mRNA (mean ± SD; n = 3). **(B)** Immunoblotting of Pum1 and Mad2 in oocytes incubated with puromycin at 0, 10, and 18 h after resumption of meiosis. **(C)** Quantitative analysis of Mad2 protein in experiments shown in C. Similar results were obtained from two independent experiments.

**Figure S2.**
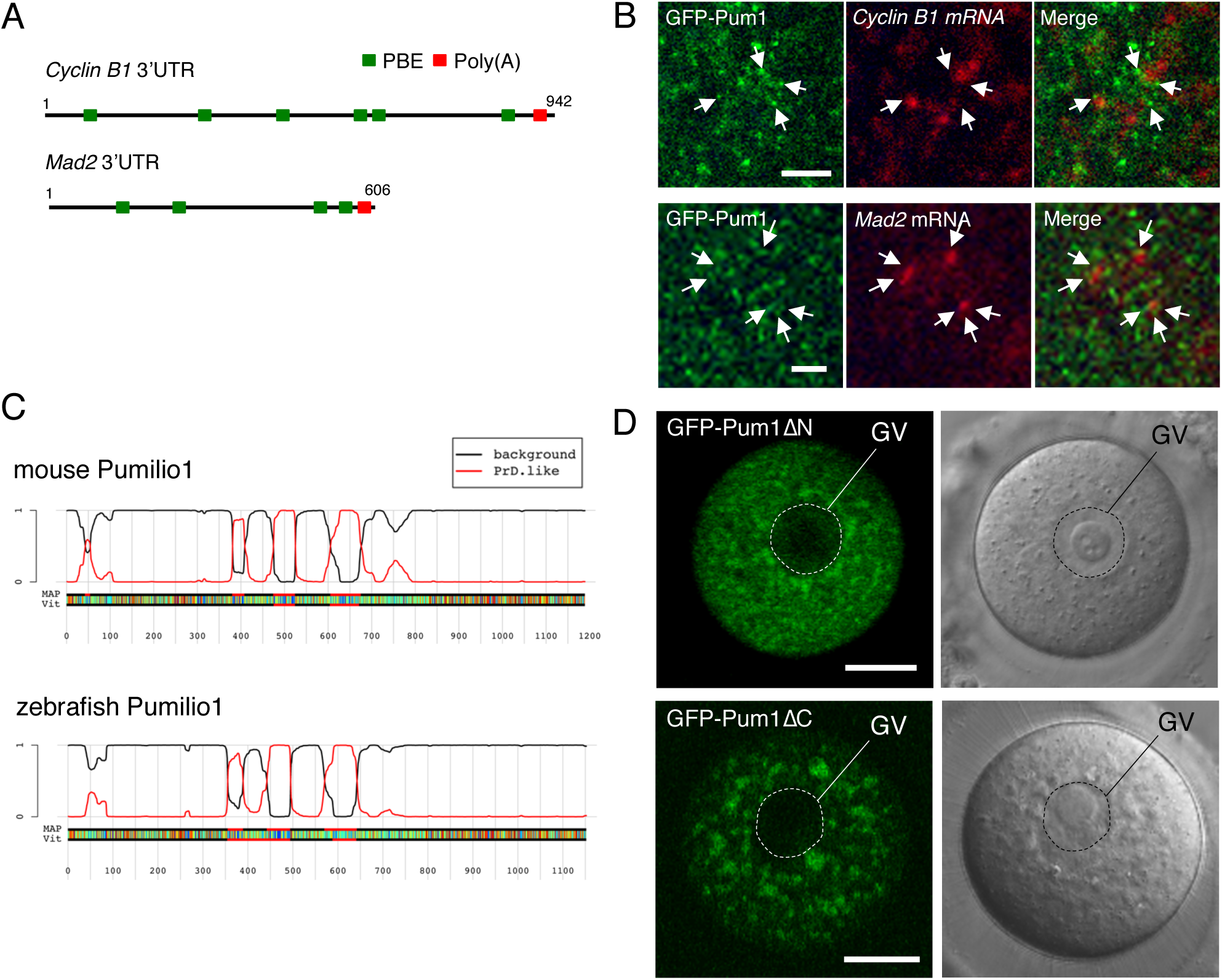
Distribution of GFP-Pum1 and that of truncated forms of Pum1. **(A)** Schematic diagrams of mouse *Cyclin B1* and *Mad2* 3’UTRs. Green rectangles indicate putative Pumilio-binding elements (PBEs), and red rectangles indicate the poly(A) signal. **(B)** FISH analysis of *Cyclin B1* (top) and *Mad2* mRNA (bottom) and immunostaining of GFP in oocytes expressing GFP-Pum1. Arrows indicate aggregates of GFP-Pum1 surrounding *Cyclin B1* or *Mad2* RNA granules. Similar results were obtained from two independent experiments. **(C)** Identification of prion-like domains by using the PLAAC web application (http://plaac.wi.mit.edu/.). **(D)** Distribution of GFP-Pum1ΔN and GFP-Pum1ΔC in immature oocytes. Similar results were obtained from six independent experiments. GV, germinal vesicle. Bar: 20 *µ*m.

**Figure S3.**
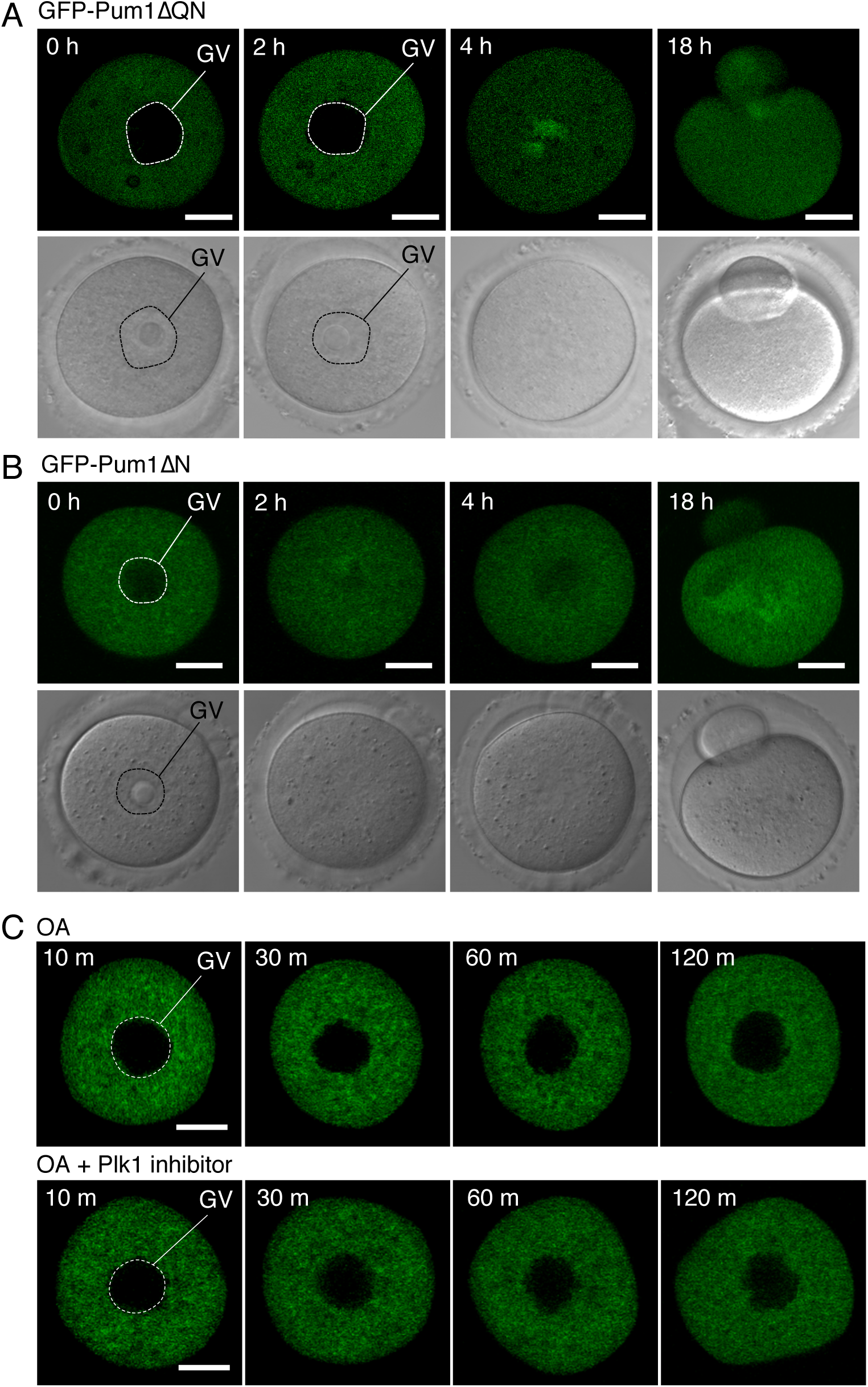
Time course of GFP-Pum1ΔQN and GFP-Pum1ΔN during oocyte maturation and that of GFP-Pum1 after OA treatment. **(A)** Time course of GFP-Pum1ΔQN at 0, 2, 4 and 18 h after resumption of meiosis. **(B)** Time course of GFP-Pum1ΔN at 0, 2, 4 and 18 h after resumption of meiosis. Similar results were obtained from two independent experiments. **(C)** Time course of GFP-Pum1 in oocytes treated with OA or OA and Plk1 inhibitor 0-120 min after treatment. Similar results were obtained in 6 oocytes from two independent experiments. GV, germinal vesicle. Bars: 20 *µ*m.

